# ImmunoCellCycle-ID: A high-precision immunofluorescence-based method for cell cycle identification

**DOI:** 10.1101/2024.08.14.607961

**Authors:** Yu-Lin Chen, Yu-Chia Chen, Aussie Suzuki

**Affiliations:** McArdle Laboratory for Cancer Research, Department of Oncology, University of Wisconsin-Madison, Madison, Wisconsin, USA; Molecular and Cellular Pharmacology Graduate Program, University of Wisconsin-Madison, Madison, Wisconsin, USA; Carbone Comprehensive Cancer Center, University of Wisconsin-Madison, Madison, Wisconsin, USA

## Abstract

The cell cycle is a fundamental process essential for cell proliferation, differentiation, and development. It consists of four major phases: G1, S, G2, and M. These phases collectively drive the reproductive cycle and are meticulously regulated by various proteins that play critical roles in both the prevention and progression of cancer. Traditional methods for studying these functions, such as flow cytometry, require a substantial number of cells to ensure accuracy. In this study, we have developed a user-friendly, immunofluorescence-based method for identifying cell cycle stages, providing single-cell resolution and precise identification of G1, early S, late S, early G2, late G2, and each sub-stage of the M phase using fluorescence microscopy. This method provides high-precision cell cycle identification and can serve as an alternative to, or in combination with, traditional flow cytometry to dissect detailed substages of the cell cycle in a variety of cell lines.

## Introduction

Cell cycle is a crucial process for proliferation, differentiation, and development in all organisms. It is precisely regulated by several checkpoint machineries that monitor and correct errors to ensure normal cell cycle progression (Harper and Brooks, 2005; Schafer, 1998; Vermeulen et al., 2003). Failures in this system often lead to carcinogenesis and tumor progression (Matthews et al., 2022). The reproductive cell cycle consists of four stages: G1, S, G2, and M phases. G1, S, and G2 phases are collectively known as interphase (Harper and Brooks, 2005; Schafer, 1998; Vermeulen et al., 2003). During G1 phase, cells increase in size and prepare to enter the S phase by expressing proteins required for DNA synthesis. Some cells, especially non-proliferating ones, may enter the G0 phase, which is outside the active cell cycle (Schafer, 1998; Vermeulen et al., 2003). Cyclin D, complexed with Cdk4/6, phosphorylates the retinoblastoma protein (Rb), promoting E2F-dependent gene expression and entry into S phase. In S phase, DNA polymerases synthesize the new DNA strand by adding nucleotides. After DNA replication is complete, cells progress to G2 phase to prepare for M phase. The transition from G2 to metaphase requires the activation of Cyclin B along with Cdk1. M phase, known as mitosis, includes five sub-stages: prophase, prometaphase, metaphase, anaphase, and telophase (Iemura et al., 2021).

The demand for single-cell accuracy and resolution significantly increases in broad research fields. This is attributed to the distinct patterns of protein and gene expression exhibited by various cell types and stages of the cell cycle. Flow cytometry is commonly used to detect and isolate cell populations at specific stages of the cell cycle, primarily by measuring relative DNA content (Darzynkiewicz and Juan, 2001; Rieger, 2022). G1 phase cells possess 2N DNA content (where N designates the haploid DNA content), while cells in G2/M phase have 4N DNA content and S phase cells fall between 2N and 4N. Since 2N and 4N populations are determined by relative DNA signal intensities, flow cytometry requires a significantly higher number of cells (>10,000 cells) to ensure accuracy (Darzynkiewicz and Juan, 2001). Additionally, distinguishing between G2 and M phases using traditional flow cytometry poses technical challenges because these cells have equal DNA content. Similarly, distinguishing substages within the M phase, between G1 and early S phase, and between late S phase and G2/M, can be challenging when relying solely on DNA content. To this end, EdU or BrdU labeling is employed to more accurately identify cells in the S phase, although this requires optimization of the duration of EdU or BrdU treatment (Bialic et al., 2022). Despite its widespread use, conventional flow cytometry is limited by its reliance on relative DNA content, which precludes high accuracy and precision at the single-cell level. Another common method is FUCCI (Fluorescent Ubiquitination-based Cell Cycle Indicator), which uses fluorescently labeled truncated Cdt1 and Geminin to distinguish between G1 and S/G2/M phases in live cell imaging or flow cytometry (Sakaue-Sawano et al., 2008; Zielke and Edgar, 2015). Although this technique is powerful for cell cycle identification, it requires the creation of stably or transiently expressing cell lines and is technically challenging for distinguishing more detailed cell cycle stages. Other fluorescence microscopy-based methods for cell cycle detection largely rely on DNA morphology and content, similar to flow cytometry, making it challenging to accurately determine detailed cell cycle stages (Bruhn et al., 2014; Roukos et al., 2015; Yamazaki et al., 2020).

In this study, we developed the ImmunoCellCycle-ID method, an immunofluorescence-based technique for identifying cell cycle stages with single-cell resolution and high accuracy. We demonstrate its effectiveness and robustness using several common cell lines. As this method employs standard immunofluorescence techniques and conventional fluorescence microscopy, it is cost-effective, user-friendly, and accessible for most researchers. This approach will be invaluable for investigating stage-specific regulatory mechanisms in the cell cycle.

## Results

### Screening cell cycle regulated proteins

To identify proteins that could potentially be used to determine cell cycle stages, we screened several major proteins known to regulate specific cell cycle stages using immunofluorescence microscopy. These proteins included Cdt1, PCNA, Cyclin B1, phospho-Histone H3 S10, CENP-F, phospho-Rb, Geminin, Cdk4, Centrin, γ-tubulin, p53, Lamin A, and α-tubulin (**Fig. 1**). Cdt1 and Geminin are used in the FUCCI cell cycle live imaging system (Sakaue-Sawano et al., 2008). As expected, Cdt1 localized to the nucleus only during the G1 phase, while Geminin began its localization to the nucleus early in the S phase and maintained its expression until anaphase. Histone H3 Ser-10, a substrate for Aurora B, is phosphorylated specifically during mitosis and is traditionally used as a mitotic marker (Hirota et al., 2005). Consistent with this, phospho-Histone H3 S10 was present from prophase and continued until metaphase, then significantly dropped to undetectable levels in anaphase. The nuclear envelope, labeled by Lamin A, remained intact until prophase when the nuclear envelope breaks down. As expected, Cdk4 was more strongly detected in the G1 phase compared to other cell cycle stages. Centrosome duplication began in the S phase and was completed in the G2 phase (see Centrin, γ-tubulin, and α-tubulin in **Fig. 1**). p53, a tumor suppressor, was found in the nucleus during G1, then formed puncta in the nucleus during S phase, likely at sites of DNA damage, and was also present in the centrosome from the G2 phase (Contadini et al., 2019; Oikawa et al., 2024). Phospho-Rb (Ser807/811) was specifically detected from early S phase to prophase (Narasimha et al., 2014; Sanidas et al., 2019). Cyclin B1 cytosolic levels were significantly increased during G2 phase with an accumulation at centrosomes. Subsequently, Cyclin B1 translocated into the nucleus during prophase, and persisted until anaphase onset (Lindqvist et al., 2007). PCNA (Proliferating Cell Nuclear Antigen), a well-documented marker for DNA synthesis, plays a crucial role in both DNA replication and repair (Schonenberger et al., 2015). PCNA was detected in the nucleus from G1 to G2 phase (details discussed in the next section). CENP-C is a component of inner kinetochore CCAN (constitutive centromere associated network) proteins (Musacchio and Desai, 2017). It localizes at kinetochores throughout the cell cycle. CENP-F, known for stabilizing kinetochore-microtubule attachments as a kinetochore corona protein, was previously proposed as a G2 phase marker because it accumulated at nucleoplasm in G2 before moving to kinetochores (Berto and Doye, 2018; Hussein and Taylor, 2002; Liao et al., 1995; Wynne and Vallee, 2018). On the contrary, our findings revealed that CENP-F entered the nucleus as early as the early S phase. Although the expression levels and cellular distribution of these selected proteins were roughly regulated based on cell cycle stages, accurately identifying all cell cycle stages using these markers alone remains challenging.

**Figure 1:**
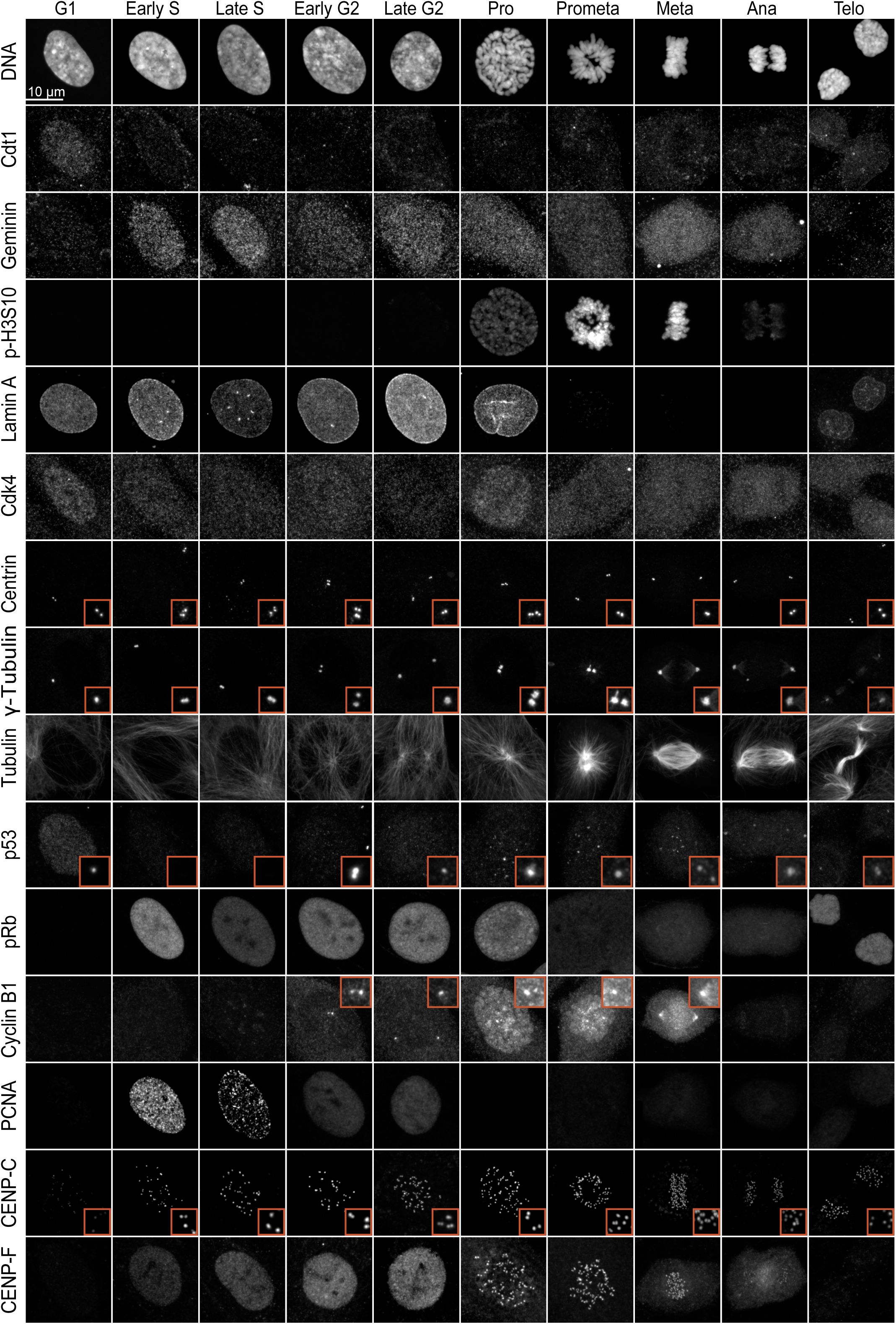
Screening the spatiotemporal regulation of key cell cycle proteins. Representative immunofluorescence images of cells labeled with Cdt1, Geminin, phospho-Histone H3 (Ser10), Lamin A, Cdk4, Centrin, γ-tubulin, α-tubulin, p53, phospho-Rb, Cyclin B1, PCNA, CENP-C, and CENP-F at different cell cycle stages.

### Immunofluorescence-based high-precision Cell Cycle IDentification (ImmunoCellCycle-ID) method

To develop a method that significantly enhances the accuracy of identifying cell cycle stages through immunofluorescence, we employed immunofluorescence labeling with a combination of selected markers: CENP-C, CENP-F, PCNA, and DNA (**Fig. 2A**). We tested specific antibodies for these proteins and detailed the staining conditions in the **Methods** section, **Supplementary Figure 1**, and **Supplementary Table 1**. Mitotic sub-stages were determined by DNA staining: prophase was characterized by chromatin condensation before nuclear envelope breakdown (NEBD), prometaphase followed NEBD without the complete formation of the metaphase plate, metaphase presented a fully aligned metaphase plate, anaphase featured the partitioning of sister chromatids, and telophase involved the reformation of the nuclear envelope while the daughter cells remained connected via the midbody (**Fig. 2A**) (McIntosh, 2016).

**Figure 2:**
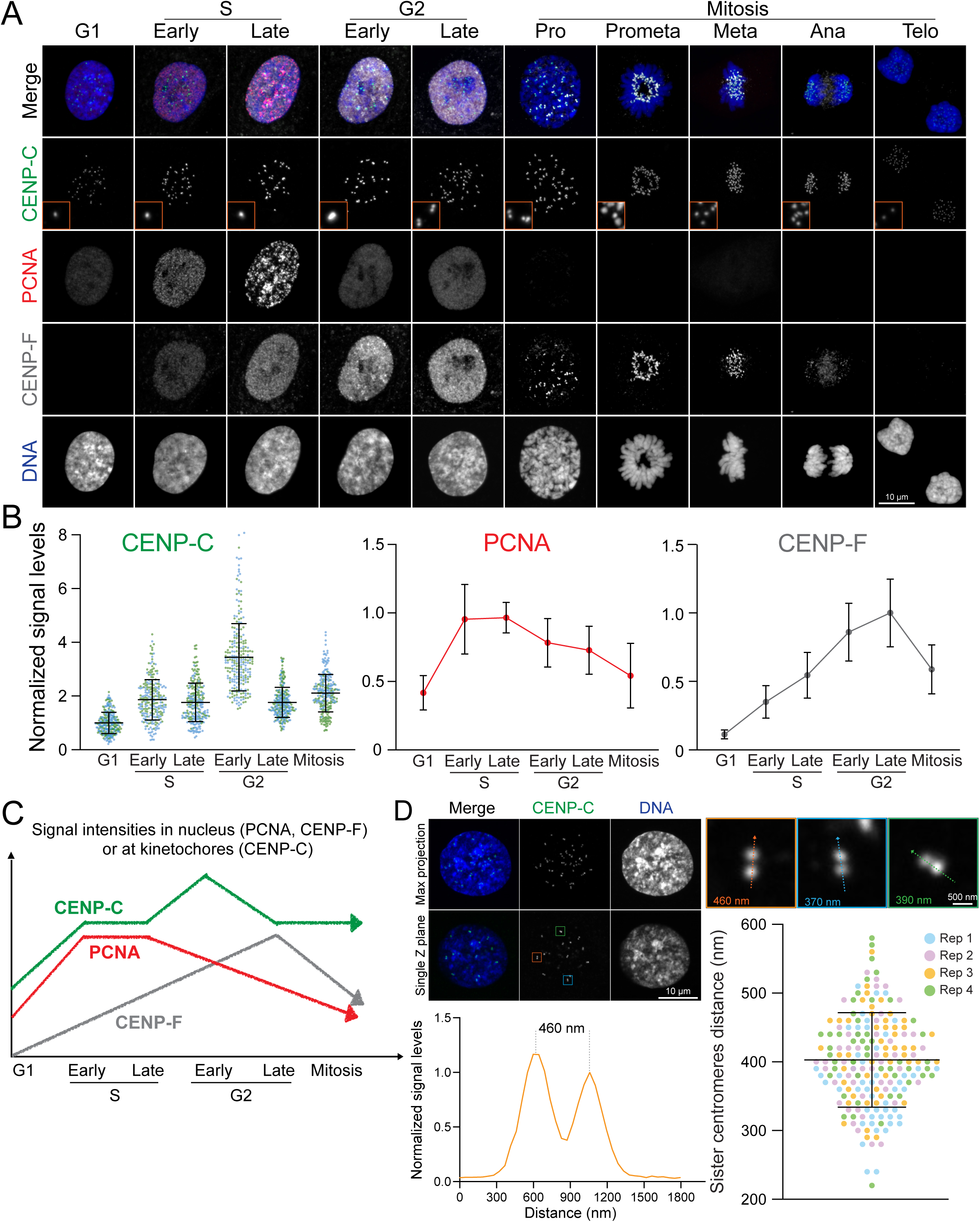
Immunofluorescence-based Identification of Cell Cycle Stages. (A) Representative immunofluorescence images of cells at each cell cycle stage, labeled with CENP-C, PCNA, and CENP-F. (B) Quantification of relative nuclear signal intensities for PCNA, CENP-F, and CENP-C at different cell cycle stages. For PCNA and CENP-F, n = 20; for CENP-C, n = 10, kinetochores = 250 (from two replicates). (C) Schematic representation illustrating the dynamics of nuclear signal variations in CENP-C, CENP-F, and PCNA as markers for identifying each cell cycle stage. (D) Measurement of the distance between sister centromeres. Top: Representative image of late G2 phase cells. Bottom left: Line scan of the distance between sister centromeres. Bottom right: Quantification of sister centromere distance in late G2 stage. n = 200 (from four replicates).

Although PCNA was detected in the nucleus throughout the entire interphase, it exhibited distinct spatial organization during the S phase. In the G1 phase, it was uniformly localized to the nucleus, but its presence significantly increased within the nucleus and appeared as small puncta across the nuclei in the early S phase, before forming more distinct and less uniform puncta in the late S phase. In the G2 phase, similar to the G1 phase, the PCNA nuclear signals decreased and became uniformly distributed (**Fig. 2A-C and S1**). Consequently, it is impossible to distinguish between G1 and G2 phases by PCNA and DNA staining alone. This observation challenges its effectiveness as a specific G2 marker. Therefore, we characterized cellular phases based on CENP-F and PCNA signals: G1 phase cells exhibited no nuclear signals for CENP-F but displayed uniform PCNA nuclear signals; early S phase cells showed CENP-F nuclear signals alongside small, brighter punctuated PCNA signals; late S phase cells were marked by distinct, bright punctuated nuclear signals of PCNA; and G2 phase cells were identified by the presence of CENP-F nuclear signals in the absence of punctuated PCNA signals.

The traditional hallmark of the G2 phase is the presence of paired kinetochore/centromere signals within the interphase nucleus (**Fig. 2A and 2D**). Kinetochores, elaborate macromolecular protein complexes situated on centromeric chromatin, act as pivotal platforms for microtubule assembly, playing a critical role in orchestrating chromosome segregation during mitosis. While numerous kinetochore proteins are specifically recruited prior to or during mitosis, the structural core of kinetochores, the Constitutive Centromere Associated Network (CCAN) proteins, remains anchored to the centromeric chromatin throughout the cell cycle (Cheeseman and Desai, 2008; Musacchio and Desai, 2017). Centromeric DNA undergoes replication during the S phase, concurrently with other DNA regions. During the G2 phase, replicated sister centromeres became separated by approximately 400 nm, which can be resolved by high-resolution fluorescence microscopy (**Fig. 2D**). In our investigation, we specifically labeled CENP-C, a major component of CCAN, as a definitive marker for centromeres/kinetochores (Musacchio and Desai, 2017). As expected, CENP-C levels at kinetochores significantly increased during S phase due to the recruitment of new CENP-C to newly synthesized centromeres (Gascoigne and Cheeseman, 2013). We also observed that all CENP-C foci became closely positioned pairs during late G2 phase (**Fig. 2A and 2D**). Intriguingly, our studies identified cells exhibiting CENP-F nuclear signals devoid of PCNA, yet these cells maintained singular CENP-C foci. Consequently, we classified these cells as the early G2 phase (**Fig. 2A-C**). Importantly, in these cells, CENP-C signal intensities reached their peak, indicating that all centromeres had completed synthesis and CENP-C had been recruited to these newly synthesized centromeres, even though the centromeres had not yet begun to separate. To detect the early G2 phase, labeling any CCAN protein is effective. However, labeling CENP-A, CENP-B, or using ACA (anti-centromere antibody) is not effective, as the amounts of these proteins do not increase at kinetochores during the early G2 phase relative to the G1 phase, unlike CCAN proteins.

### Percentages of a cell population in the different phases of the cell cycle in RPE1 cells

We next assessed the distribution of cell cycle phases in asynchronous RPE1 cells using ImmunoCellCycle-ID method (**Fig 3A-C and S3**). Cells at the logarithmic growth phase, achieving approximately 60-70% confluency, were fixed and subsequently stained (see **Methods**). Our findings revealed that about 49% of the cells were in the G1 phase, 27% in the S phase, 18% in the G2 phase, and 6% were undergoing mitosis (**Fig 3B**). Flow cytometry analysis from this and previous studies using RPE1 cells showed a range of 52-64% in the G1 phase, 15-21% in the S phase, and 18-24% in the G2/M phase, aligning with the ImmunoCellCycle-ID analysis (**Fig. 3D**) (Lau et al., 2009; McKinley and Cheeseman, 2017; Pei et al., 2022). Although cells with 4N DNA content (G2/M phases) constituted around 24% of asynchronous RPE1 cells in ImmunoCellCycle-ID analysis, the majority were in the G2 phase rather than in mitosis, with approximately 75% of the G2/M population in G2 phase (**Fig. 3B**). Additionally, within S phase cells, approximately 78% were in early S phase, and 22% were in late S phase (**Fig. 3B**). Similarly, within G2 phase cells, a predominance of cells was in early G2 phase (∼72%) over late G2 phase (∼28%), diverging from the conventional classification. This discrepancy could account for the observed differences in G2 phase frequencies between flow cytometry and fluorescence microscopy, with the latter often observing a lower G2 cell population than anticipated from flow cytometry. We also quantified the frequency of each sub-stage within mitosis. The majority of cells were in telophase and metaphase, constituting approximately 40% and 30%, respectively. The remaining sub-stages were nearly equal with each comprising around 10% (**Fig. 3C**). Through our study, we have not only demonstrated but also validated the accuracy and reliability of our immunofluorescence-based technique in precisely delineating the stages of the cell cycle.

**Figure 3:**
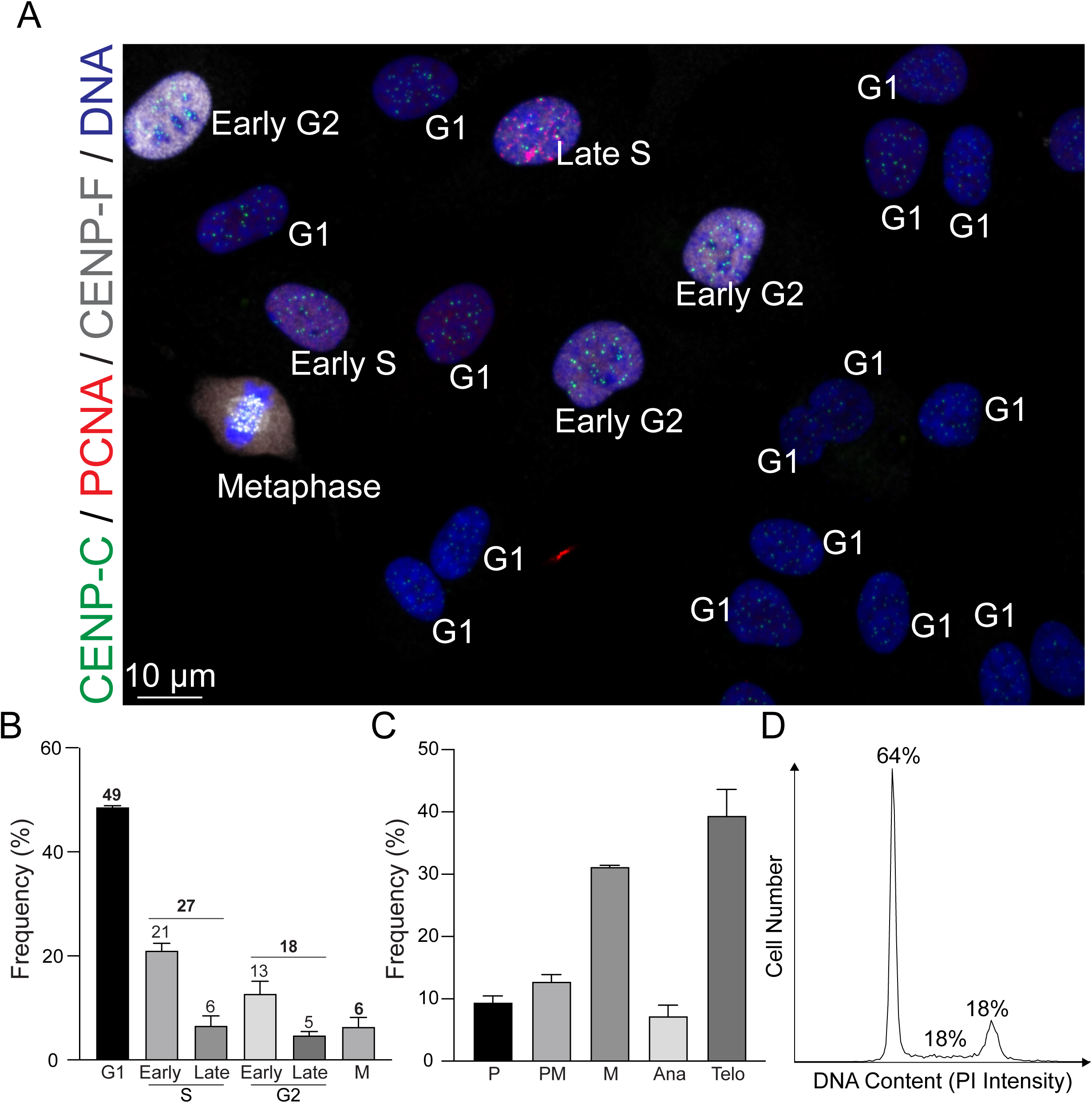
Analysis of Cell Cycle Distribution in Asynchronous RPE1 Cells. (A) Representative immunofluorescence images of asynchronous RPE1 cells labeled with CENP-C, PCNA, and CENP-F. (B) Distribution of cells across different phases of the cell cycle. n = 424 (from two replicates). (C) Proportion of mitotic cells within each sub-stage of mitosis. n = 302 (from two replicates). (D) Representative flow cytometry histograms showing detection of DNA content (PI signal intensity) in RPE1 cells.

### Performance of ImmunoCellCycle-ID Across Various Cell Types

To demonstrate the robustness of ImmunoCellCycle-ID in cell cycle stage determination is not limited to non-transformed cell lines, we applied our method to different cancer cell lines, including Cal51 (a triple-negative breast cancer cell line), HCT116 (a colon cancer cell line), HeLa (a cervical cancer cell line), T47D (a luminal A subtype breast cancer cell line), and U2OS (an osteosarcoma cell line) (**Fig. 4A**). As expected, we accurately determined each stage of the cell cycle in all cell types without altering the fixation, staining, and imaging protocols. All the selected cancer cell lines exhibited 48-67% of cells in the G1 phase. Except for T47D, the other four cell lines showed slightly higher populations in the S phase compared to RPE1 cells. In RPE1, Cal51, and U2OS cells, the majority of the S phase was in the early S phase, whereas HCT116, T47D, and HeLa cells exhibited either a similar distribution between early and late S phases or a higher population in the late S phase. Additionally, we demonstrated that ImmunoCellCycle-ID is capable of identifying all sub-stages of mitosis in all cell types (**Fig. 4A, left**). To validate the ImmunoCellCycle-ID results, we also performed flow cytometry analysis on selected cell lines (Cal51, HCT116, and HeLa) (**Fig. 4B**). These flow cytometry results were consistent with the measurements obtained by ImmunoCellCycle-ID methods. These results confirm the accuracy and reliability of ImmunoCellCycle-ID in determining cell cycle stages and populations.

**Figure 4:**
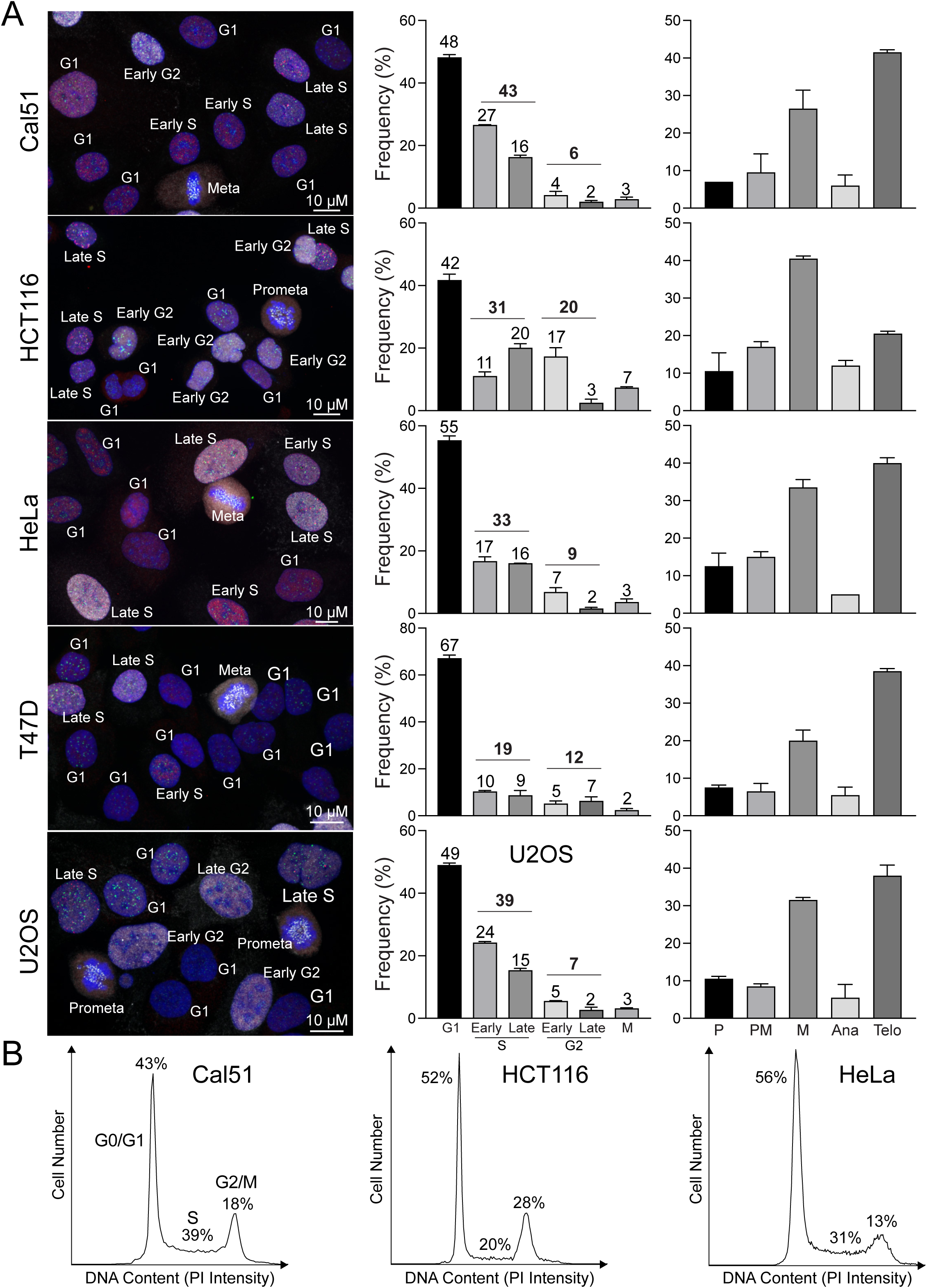
Cell Cycle Distribution in various cancer cell lines. (A) Left panel: representative immunofluorescence images of asynchronous Cal51, HCT116, HeLa, T47D, and U2OS cells labeled with CENP-C, PCNA, and CENP-F. Middle panel: distribution of cells across different phases of the cell cycle. From top to bottom, n = 455, 447, 518, 467, and 437 (from two replicates). Right panel: proportion of mitotic cells within each sub-stage of mitosis. From top to bottem, n = 181, 201, 212, 156, and 188. (from two replicates). (B) Representative flow cytometry histograms showing detection of DNA content (PI signal intensity) in Cal51, HCT116, and HeLa cells.

### Limitation of this study

Our immunofluorescence-based cell cycle identification method provides single-cell accuracy, is accessible, and user-friendly; however, it requires cell fixation and cannot be used with live cells. Additionally, mitotic cells tend to detach more easily compared to cells in other stages of the cell cycle, necessitating gentle fixation to minimize underestimation of the mitotic population. While we demonstrated this method using adherent cell lines, it can also be applied to floating cells using the cytospin or other alternative methods.

## Discussion

Determining the populations at each stage of the cell cycle is a common experiment across a broad range of research fields. Traditionally, this is performed using flow cytometry (Pozarowski and Darzynkiewicz, 2004; Rieger, 2022), which provides accurate results based on the number of cells measured. However, conventional flow cytometry lacks single-cell accuracy and cannot distinguish between G2 and the sub-stages of mitosis, cells at the borders of cell cycle stages, or between G2/M and cells with whole-genome duplications (Banfalvi, 2011; Darzynkiewicz and Juan, 2001; Rieger, 2022). Our immunofluorescence-based cell cycle identification method offers a user-friendly fluorescence microscopy approach with single-cell accuracy. It can be used not only for population analysis, but also for a single cell cell-cycle determination. The advantages of this method include its capacity to precisely identify G1, early S, late S, early G2, late G2, and every sub-stage of mitosis (**Fig. 2**). In this study, we demonstrate and validate the accuracy and robustness of this method and define new sub-stages in the G2 phase, termed early and late G2 phases.

Flow cytometry is a widely-used technique for cell cycle analysis but often requires access to an institutional flow cytometry core. In contrast, the ImmunoCellCycle-ID method only necessitates the use of conventional fluorescence microscopy, which is now standard equipment in many laboratories. Flow cytometry faces technical challenges in accurately distinguishing the boundaries between the end of G1-phase and the beginning of S-phase, as well as the end of S-phase and G2/M phase. Additionally, aneuploid cells and chromosomally unstable cells, such as many cancer cells, are more difficult to analyze accurately using conventional flow cytometry. On the other hand, ImmunoCellCycle-ID utilizes homeostasis protein markers that are spatiotemporally regulated in their cellular dynamics across different cell types, including non-transformed and transformed cells with various karyotypes. This method provides reproducible and high-precision cell cycle identification regardless of cell type. In summary, by employing standard immunofluorescence techniques and conventional fluorescence microscopy, the ImmunoCellCycle-ID method is a useful, cost-effective, and accessible tool for researchers investigating stage-specific regulatory mechanisms in the cell cycle.

## Acknowledgement

We would like to thank Yoshitaka Sekizawa and Yokogawa Electric Corporation for critical equipment and technical support. We would also like to thank Dr. Stephen Taylor for generously providing the antibodies for CENP-F, and Syon Reddy for his support in quantification. Part of this work is supported by Wisconsin Partnership Program, Research Forward from the University of Wisconsin-Madison Office of the Vice Chancellor for Research with funding from the Wisconsin Alumni Research Foundation, start-up funding from University of Wisconsin-Madison SMPH, UW Carbone Cancer Center, and McArdle Laboratory for Cancer Research, and NIH grant R35GM147525 (to A.S.).

## Author contribution

YL.C. conducted precision imaging experiments and analyses. YC.C established the immunofluorescence-based cell cycle identification method. A.S. conceptualized, supervised, and funded the project. A.S. prepared the initial manuscript draft. All authors reviewed and contributed to the manuscript’s refinement.

## Competing Financial Interests

The authors declare no further conflict of interests.

## Methods

### Cell Culture

All cell lines were originally obtained from the American Type Culture Collection (ATCC, Manassas, VA, USA). RPE1, Cal51, HeLa, T47D, U2OS, and HCT116 cells were grown in DMEM high glucose (Cytiva Hyclone; SH 30243.01) supplemented with 1% penicillin-streptomycin, 1% L-glutamine, and 10% fetal bovine serum under 5% CO_2_ at 37°C in an incubator.

### Immunofluorescence

RPE1 cells were fixed by 4% PFA (Sigma) or 100% Methanol. Cells which fixed with PFA were then permeabilized by 0.5% NP40 (Sigma) and incubated with 0.1% BSA (Sigma). Primary and secondary antibodies are listed in Supplementary Table 1. Stained samples were imaged by CSU W1 SoRa spinning disc confocal, which was equipped with Uniformizer and a Nikon Ti2 inverted microscope with a Hamamatsu Flash V2 camera and a 100x Oil objective (NA = 1.40). Microscope system was controlled by Nikon Elements software (Nikon).

### Image analysis

Image analysis was performed using Nikon Elements software (Nikon) or Metamorph (Molecular Devices). For signal quantification in nucleus or kinetochores, we utilized local background correction methods used in previous study (Suzuki et al., 2015). Intra-kinetochore distances in late G2 phase were obtained by measuring the distance between the peaks of signal intensity (Loi et al., 2023)

### Flow cytometry

RPE1, HeLa, Cal51, and Hct116 were fixed in 70% cold ethanol at 4°C for 3 hours. DNA staining was performed with 100 µg/mL RNase A (Sigma), 25 µg/mL Propidium Iodide (Sigma), and 0.1% Triton X-100 (Sigma) at 4°C for 18 hours. Analysis was performed on a ThermoFisher Attune NxT cytometer with Attune software. Cell-cycle modeling was performed with ModFit 5 software. Cells were gating by the PI signal area to identify signal cells for analysis.

### Statistics

All experiments were independently repeated 2-3 times for mitotic duration measurements. p-values were calculated using one-way ANOVA and the two-tailed Student’s t-test. p-values <0.05 were considered significant.

## Notes

### Competing Interest Statement

The authors have declared no competing interest.

